# Streamlined Magnetic Resonance Fingerprinting: Fast Whole-brain Coverage with Deep-learning Based Parameter Estimation

**DOI:** 10.1101/2020.11.28.400846

**Authors:** Mahdi Khajehim, Thomas Christen, Fred Tam, Simon J. Graham

**Affiliations:** Department of Medical Biophysics, University of Toronto, Toronto, ON, Canada; Grenoble Institute of Neuroscience, Inserm, Grenoble, France; Hurvitz Brain Sciences Research Program, Sunnybrook Research Institute, Toronto, ON, Canada

**Keywords:** quantitative MRI, MR fingerprinting, echo-planar imaging (EPI), combined gradient echo and spin echo, T1 mapping, T2 mapping, T2* mapping

## Abstract

Magnetic resonance fingerprinting (MRF) is a novel quantitative MRI (qMRI) framework that provides simultaneous estimates of multiple relaxation parameters as well as metrics of field inhomogeneity in a single acquisition. However, current bottlenecks exist in the forms of (1) scan time; (2) need for custom image reconstruction; (3) large dictionary sizes; (4) long dictionary-matching time. The aim of this study is to introduce a novel streamlined magnetic-resonance fingerprinting (sMRF) framework that is based on a single-shot echo-planar imaging (EPI) sequence to simultaneously estimate tissue T1, T2, and T2* with integrated B1^+^ correction. Encouraged by recent work on EPI-based MRF, we developed a method that combines spin-echo EPI with gradient-echo EPI to achieve T2 in addition to T1 and T2* quantification. To this design, we add simultaneous multi-slice (SMS) acceleration to enable full-brain coverage in a few minutes. Moreover, in the parameter-estimation step, we use deep learning to train a deep neural network (DNN) to accelerate the estimation process by orders of magnitude. Notably, due to the high image quality of the EPI scans, the training process can rely simply on Bloch-simulated data. The DNN also removes the need for storing large dictionaries. Phantom scans along with in-vivo multi-slice scans from seven healthy volunteers were acquired with resolutions of 1.1×1.1×3 mm^3^ and 1.7×1.7×3 mm^3^, and the results were validated against ground truth measurements. Excellent correspondence was found between our T1, T2, and T2* estimates and results obtained from standard approaches. In the phantom scan, a strong linear relationship (R=1-1.04, R^2^>0.96) was found for all parameter estimates, with a particularly high agreement for T2 estimation (R^2^>0.99). Similar findings are reported for the in-vivo human data for all of our parameter estimates. Incorporation of DNN results in a reduction of parameter estimation time on the order of 1000 x and a reduction in storage requirements on the order of 2500 x while achieving highly similar results as conventional dictionary matching (%differences of 7.4±0.4%, 3.6±0.3% and 6.0±0.4% error in T1, T2, and T2* estimation). Thus, sMRF has the potential to be the method of choice for future MRF studies by providing ease of implementation, fast whole-brain coverage, and ultra-fast T1/T2/T2* estimation.

## 1. Introduction

Quantitative magnetic resonance imaging (qMRI) typically refers to the quantitative mapping of tissue parameters such as T1 and T2 (Cheng et al., 2012; Deoni, 2010; Keenan et al., 2019; Tofts, 2005). Compared to the clinically dominant qualitative (e.g. T1- and T2-weighted) techniques, qMRI provides an important, objective alternative in clinical and research settings for effective detection and monitoring of different neurological pathologies (Cheng et al., 2012), including stroke (Bernarding et al., 2000), neurodegenerative diseases (Baudrexel et al., 2010; Rugg-Gunn et al., 2005; Townsend et al., 2004) and brain tumors (Just and Thelen, 1988; Kurki and Komu, 1995). However, qMRI is typically limited by very long acquisition times, constituting a barrier for routine clinical practice (Cheng et al., 2012; Keenan et al., 2019; Tofts, 2005; Whittall et al., 1997). Moreover, conventional methods suffer from high sensitivity to scanner imperfections (e.g. B1^+^ or B0 inhomogeneities), such that T1 and T2 estimates from different scanners and different methods may not be directly comparable (Stikov et al., 2015). As a result, there is a clear need for faster and more robust quantitative imaging approaches that can simultaneously estimate conventional tissue parameters (T1, T2, and T2*) as well as field-specific parameters (B0 and B1^+^).

Accelerated qMRI methods have been proposed to tackle certain parameters (Cunningham et al., 2006; Deoni et al., 2004; Ehses et al., 2013; Freeman et al., 1998; Heule et al., 2014), but most of them are incapable of estimating more than two tissue parameters at once. Moreover, these methods sometimes rely on separate B1 field mapping scans to increase accuracy. To meet these challenges, magnetic resonance fingerprinting (MRF) was proposed as a novel qMRI framework that provides simultaneous estimates of multiple relaxation parameters as well as metrics of field uniformity in a single acquisition (Cloos et al., 2016; Ma et al., 2013). In MRF, sequence parameters are varied dynamically in a pseudorandom fashion and the acquired signal time courses are matched to a pre-calculated dictionary (created using Bloch simulations) through a pattern-matching algorithm. The matching identifies the dictionary entry and its corresponding set of predetermined qMRI parameters. So far, MRF has mostly provided quantification of T1 and T2 relaxation times and is most commonly based on a spiral readout with a large undersampling factor to speed up acquisition (Bipin Mehta et al., 2018; Jiang et al., 2014; Ma et al., 2013; Poorman et al., 2019). The spiral readouts were chosen for their high sampling efficiency, and are rotated at each TR to randomize undersampling artifacts and to provide spatial incoherence (Bipin Mehta et al., 2018). Despite MRF’s great promise, there remain several bottlenecks:

1. Spiral acquisitions usually require an offline image reconstruction, including regridding and sampling density compensation and are more vulnerable to field imperfections as compared to Cartesian readouts. Based on the latter, MRF using echo-planar imaging (EPI) was introduced recently (Hermann et al., 2020; Rieger et al., 2018, 2017), but so far can only provide T1 and T2* estimates.
2. The original spiral-based MRF framework has the opposite issue --- it was designed for T1 and T2 quantification but not T2*. The current spiral-based approaches that offer whole-brain simultaneous T1, T2 and T2* estimation (Hong et al., 2018; C. Y. Wang et al., 2019; Wyatt et al., 2018) require relatively lengthy acquisitions (typically about 1/2 minute per slice) by clinical standards.
3. In the pattern matching step, the MRF estimates are quantized and the dictionary size grows very rapidly with the increasing number of parameters (dictionary dimensionality) and steps, easily reaching hundreds of gigabytes. This constitutes another barrier to the broad application of MRF (Hilbert et al., 2019; Hong et al., 2018; Körzdörfer et al., 2018; Wright et al., 2018). Recently, compressing the dictionary along the time dimension and utilization of the polynomial fitting on the parameter space have been suggested as possible ways to partly circumvent this issue. Yet, due to the growing dictionary size and dimensions, more effective approaches are still required (McGivney et al., 2014; Yang et al., 2018).
4. The time needed for parameter matching is proportional to the total number of dictionary elements and depending on the hardware, can take up to several hours per slice (Hong et al., 2018). Recently, accelerated matching has been proposed (Cauley et al., 2015), but as it still relies on pattern matching, the increase in parameter estimation speed cannot fully resolve the speed bottleneck in cases of high dictionary size and dimensionality.

To address these limitations, we introduce a streamlined MRF (sMRF) framework that aims to facilitate the implementation and application of MRF by addressing the above-mentioned challenges.

1. To facilitate the image acquisition, we replace the undersampled spiral readout with a single-shot EPI readout, whereby in-plane acceleration is used to minimize image distortions. EPI-based acquisitions in MRF have been proposed previously (Rieger et al., 2017; Su et al., 2017), and allow for easy online reconstruction of images that are free from major undersampling artifacts.
2. To overcome limitations in current EPI-based MRF, we introduce a modular extension based on spin-echo (SE) EPI that provides T2 sensitivity with B1^+^ correction. In this way, two different EPI stages of the technique work in sequence to estimate T1, T2* and then T2. We further capitalize on the blipped-CAIPI simultaneous multi-slice (SMS) method (Setsompop et al., 2012; Ye et al., 2016) to achieve fast whole-brain coverage (~ 3 minutes).
3. To eliminate the need for using a large dictionary, we utilize a deep neural network (DNN) (Cohen et al., 2018; Hoppe et al., 2017) for parameter estimation that can be sufficiently trained with simulated data alone. As a result of having the regression layer as the output, this DNN-based parameter estimation stage provides non-quantized estimates and requires much lower storage than a conventional dictionary (up to orders of magnitude lower).
4. The use of DNN also allows for parameter estimation that is orders of magnitude faster than conventional dictionary matching, taking only a few minutes to compute whole-brain T1, T2, and T2* maps on a typical desktop computer (single-core, 1.6 GHz CPU).

Overall, this study is aimed at presenting an sMRF framework that is easy to implement, is fast in the data acquisition step as well as data reconstruction stage and is capable of producing estimates for all three main relaxation parameters of interest (T1, T2 and T2*) within a few minutes. We demonstrate the accuracy and robustness of the streamlined MRF technique in phantoms and in vivo human brains.

## 2. Methods

A schematic overview of the streamlined MRF pipeline is shown in Figure 1.

**Figure 1.**
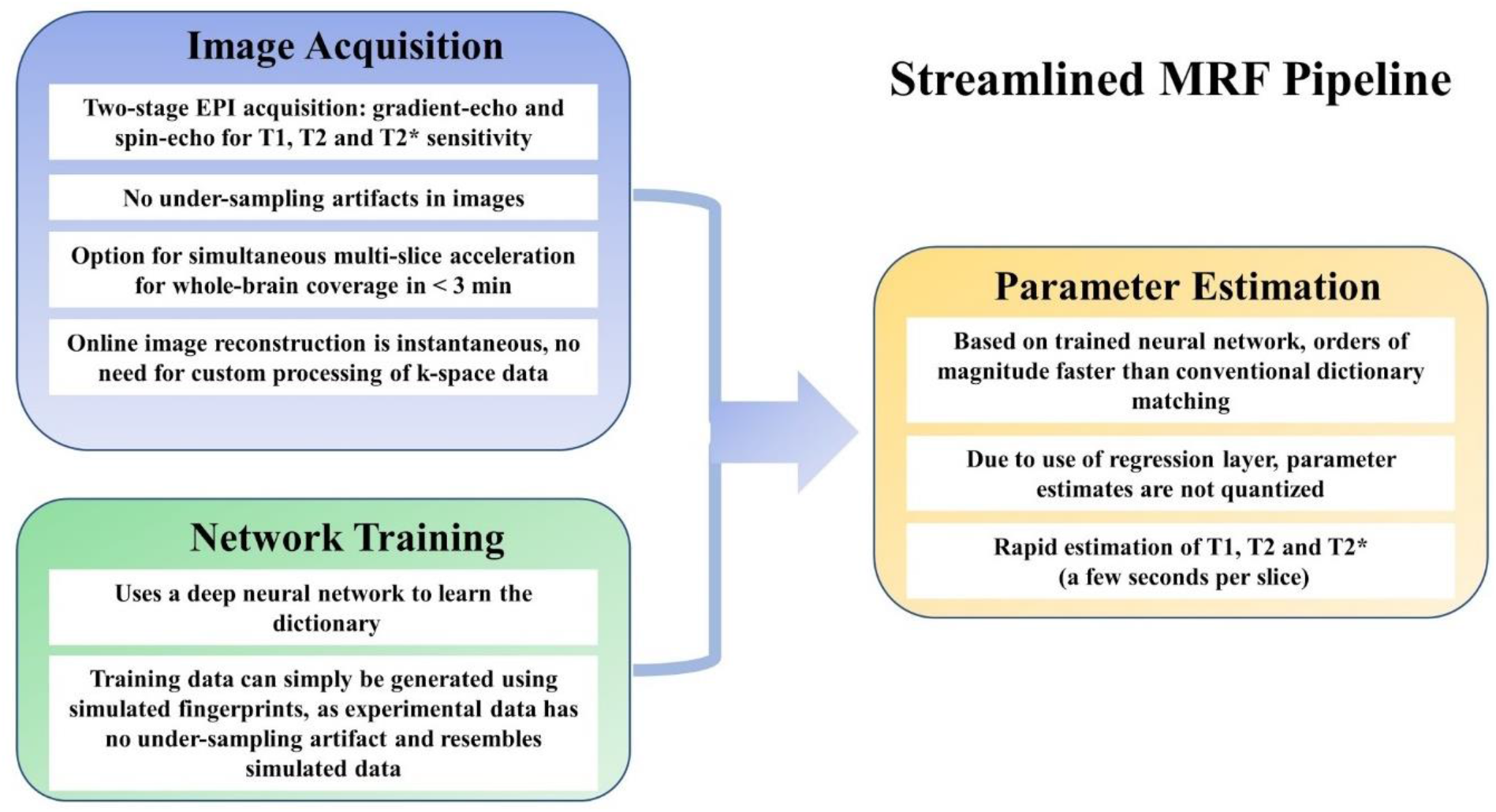
Schematic of the streamlined MRF pipeline. Our streamlined MRF method has 3 main components. *Image acquisition*, a new pulse sequence is designed that concatenates gradient- and spin-echo EPI for T1, T2 and T2* sensitivity. In the case of both, imaging parameters, including TR, TE and FA, are pseudo-randomly varied, and the images are reconstructed online, without substantial undersampling artifacts. *Network training:* as we use deep learning to replace the dictionary, a deep neural network is trained using simulated fingerprints alone, made possible due to its close resemblance to experimental data that is devoid of major undersampling artifacts. This feature greatly extends the flexibility for training. *Parameter estimation*. This is performed using the same deep neural network, and is orders of magnitude faster than pattern-matching based estimation.

### 2.1. Addressing EPI-related challenges for T2 and T2* estimation

As mentioned previously, when using EPI readouts, a unique challenge is to separate T2 from T2* effects. Our solution is to separate T2* and T2 estimation into two different stages of the sequence. The first (GE) stage builds on the recent success of EPI-based MRF to provide B1^+^ corrected T1 and T2* estimation (Rieger et al., 2018, 2017), while the second stage focuses on providing T2 estimation by using the SE contrast mechanism. In this dual-stage manner, we expect to reduce the cross-talk between T2* and T2 estimation and consequently the possibility for error propagation between these two. Moreover, another challenge in MRF is the multi-dimensional nature of the dictionary, which can easily expand storage requirements to challenging levels and result in lengthy matching processes (reportedly up to several hours per slice (Hong et al., 2018; Zhang et al., 2020)). In this respect, our two-stage approach serves as a proof of concept for a modularized design for MRF, in which each module can be optimized in parameters and duration to the estimate of interest. In this way, we constrain the complexity of the dictionary-matching process, leading to potential gains in the acquisition and computational efficiency. These outcomes will be demonstrated in the following.

### 2.2. Pulse sequence design

Our sequence design derives from the work by Rieger et al. (Rieger et al., 2018, 2017), but with a major extension. A schematic view of the sequence is shown in Figure 2. We acquire both gradient-echo (GE) and spin-echo (SE) EPI data in the same acquisition. This dual-stage design was inspired by recent work (Hong et al., 2018; C. Y. Wang et al., 2019) and motivated by the following considerations. First, as T1 and B1^+^ can both be quantified in the GE-EPI segment and then fed into the dictionary-matching process for the SE-EPI segment, it was found that even as few as 80 frames, less than half the number of frames in the GE segment (which was 200), were enough for accurate T2 estimation (see Appendix). Secondly, T2 and T2* estimations are performed from data acquired in separate (GE and SE) segments of the sequence, such that error in one estimate does not necessarily contaminate the other estimate. Lastly and importantly, such a design is advantageous over acquiring the SE-EPI and GE-EPI scans separately, as it minimizes confounding effects such as movement, time-dependent B0 inhomogeneity and possible systematic differences in auto-tuning.

**Figure 2.**
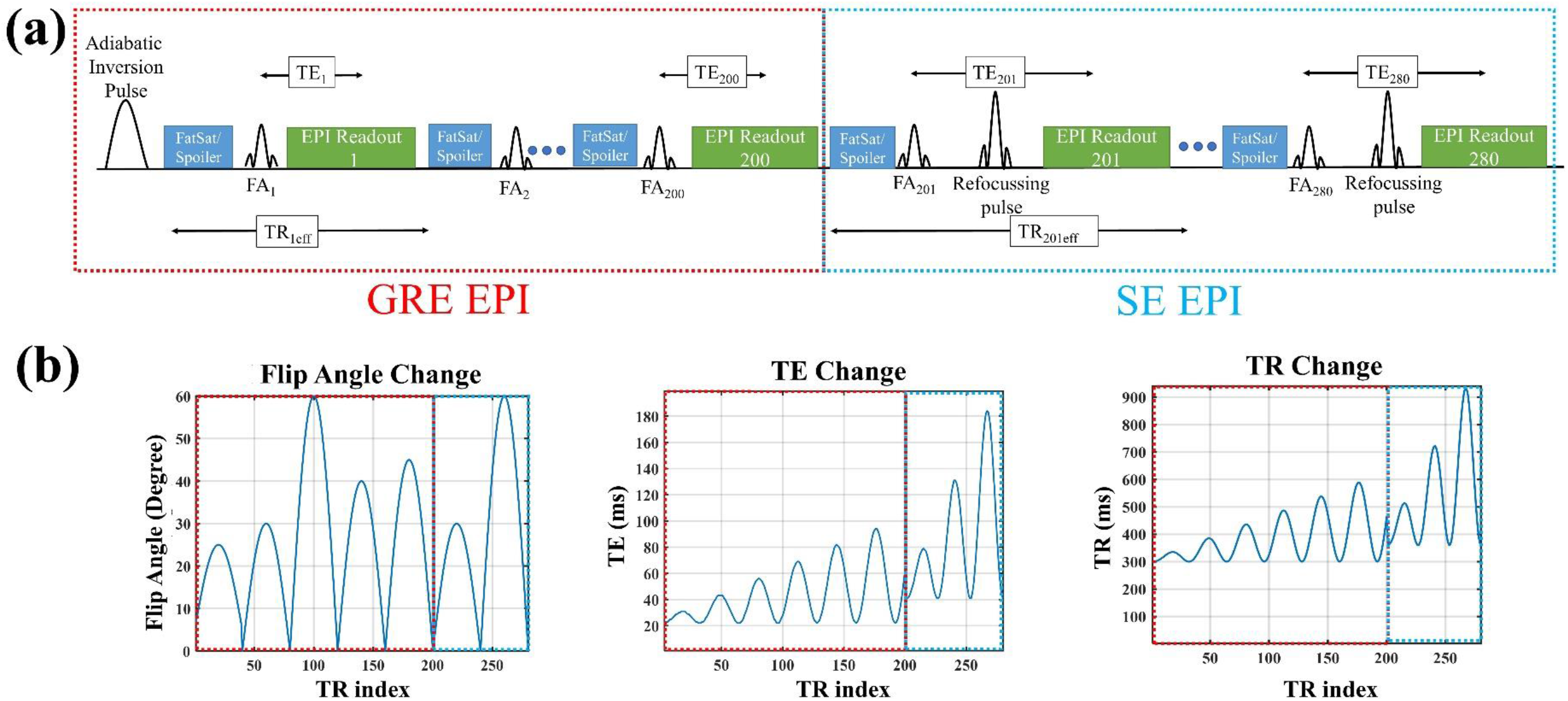
Schematic view of the sequence. (a) a Gradient Echo (GE) echo-planar imaging (EPI) stage is followed by a spin-echo (SE) stage. (b,c,d) The patterns of flip angle change, TE change and TR change, spanning the GE and SE stages (for a total of 280 frames). The red and blue dotted lines represent the GE and SE sections, respectively.

More specifically, the GE-EPI segment is similar to that proposed by Rieger et al. (Rieger et al., 2017), with only slight modifications (e.g. 200 instead of 160 frames). After a non-selective hyperbolic secant adiabatic inversion pulse, the GE-EPI segment begins (Fig. 2a): (1) the FA varies in accordance with five half periods of an amplitude-modulated sinusoidal variation (Fig. 2b), with FAs ranging from 0 to 60°; (2) TEs vary between 22-95 ms (Fig. 2c) while TR is the shortest possible for each TE as allowed by the slice-interleaved multi-slice acquisition (Fig. 2d, TR range = 300-920 ms); (3) in addition to fat saturation, both gradient and RF spoiling are implemented using crusher gradients before and after the fat saturation module (in x, y and z directions). After 200 GE-EPI frames, the sequence transitions into SE-EPI for 80 frames. The number of frames was informed by simulation (see Appendix and Supplementary Figures). Crusher gradients are added before and after the refocusing pulse in all three directions to spoil the free induction decay that may originate from non-ideal refocusing. In the SE-EPI segment, two half-periods of a sinusoidal function are used for FA variation (with the sinusoidal maxima of 30 and 60 degrees, respectively). The TE range for the SE portion is 40-190 ms. Other sequence parameters common to both GE and SE segments are: matrix size = 192×192, field of view (FOV) = 220 x 220 mm, voxel size = 1.1×1.1×3 mm, slice gap=50%, parallel imaging with GRAPPA factor=3, #reference lines=96, partial Fourier=6/8, readout bandwidth (BW)/voxel=1302 Hz, total acquisition time per slice is 29 s. Each acquisition affords multi-slice (non-whole-brain) coverage and was achieved through a standard interleaved acquisition that also has the benefit of increasing the baseline signal by lengthening the effective TR.

To enable accelerated whole-brain coverage, we further incorporated simultaneous multislice (SMS) acceleration. In the SMS implementation, we implemented blipped-CAIPI by replacing the RF excitation pulses with their SMS equivalents (excepting the initial inversion pulse, which is non-selective). The sequence parameters used for the SMS version are as follows: SMS-factor = 4, FOV shift factor = 2, matrix size =128×128, FOV= 220 x 220 mm, voxel size = 1.7×1.7×3 mm, slice gap=50%, parallel imaging with GRAPPA acceleration factor = 2, #reference lines = 24, partial Fourier = 6/8, BW/Pixel = 1954 Hz, number of slices = 24, effective acquisition time = 7 sec/slice. In-plane acceleration of at least a factor of 2 was required to minimize T2*-related image distortions.

### 2.3. Network training: Dictionary generation

In this work, as was done by Rieger et al. (Rieger et al., 2017), each voxel is represented by one isochromat. The dictionary is generated using the discrete matrix form of the Bloch equations. The effect of RF pulses is assumed to be instantaneous, and the RF-slice profile is assumed to be ideal for simplicity. B1^+^ inhomogeneity is explicitly included in the model as a scaling factor applied to the nominal FAs. By showing the range as [min:step-size:max], the simulated ranges are: T1= [100:20:4000] ms, both T2 and T2* = [5:5:30 32:2:130, 135:5:200, 210:10:350] ms and the relative B1^+^ = [0.5:0.05:1.5]. The T1, T2, and T2* values are chosen to specifically target the brain-tissue range. Overall the dictionary has ~ 350,000 entries. Dictionary generation takes less than 15 minutes on a desktop computer running on a 1.6 GHz CPU.

### 2.4. Network training: Deep learning for parameter estimation

To reduce the storage requirement and increase the parameter estimation speed in streamlined MRF, a deep learning approach was used. This stage was inspired by Cohen et al. (Cohen et al., 2018) but with some changes in the network architecture, training algorithm, the number of nodes and the number of outputs. A deep neural network (DNN) with two hidden layers was designed (consisting of 128 and 64 neurons, respectively) in which the input layer gets the measured fingerprint time series for each voxel and the output layer produces B1^+^ corrected T1 and T2* estimates. The rectified linear unit (ReLu) function was used after each hidden layer. The adaptive moment estimation (Adam) optimizer was utilized to train the network using a constant learning rate of 0.001 and the minimum batch size of 1024. 100 epochs were used for training, which took 15-20 min on a single CPU as described earlier. During the DNN training process, Gaussian white noise was added to each dictionary entry with a SNR ranging 16-40 dB. The DNN was implemented using MATLAB’s deep learning toolbox (version 12.1, Mathworks, Natick, MA, USA) using custom-written codes. Figure 3 shows a schematic view of the parameter estimation stage using DNN. The training stage produces the weights for the neurons (nodes) in the hidden layers. Note that we do not need to incorporate spatial information, and each voxel’s fingerprint is matched independently from those of others.

**Figure 3.**
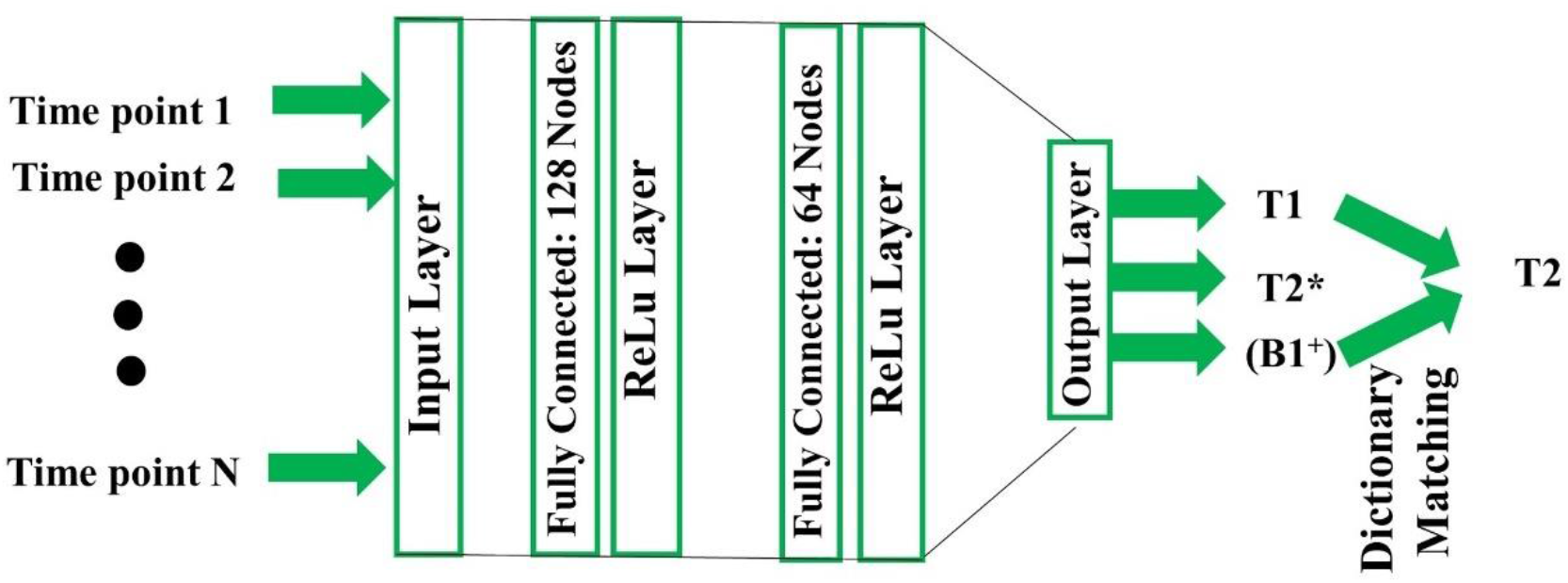
Schematic view of the DNN. The network consists of two fully connected hidden layers, with 128 nodes (neurons) and 64 nodes, respectively. The output layer produces T1 and T2* (as well as B1^+^) estimates using the GE-EPI data. The T1 and B1^+^ estimates then feed into the estimation of T2 based on the SE-EPI data. Note that the T2 estimation is performed by matching to a mini-dictionary generated using DNN-based T1 and B1^+^ results. Also, note that due to the use of the regression layer the T1 and B1^+^ estimates are continuous, although the dictionary used in the training process was based on quantized relaxation values.

As our acquired images and hence the MRF time series are free from major under-sampling artifacts, we have the advantage of requiring simply the simulated dictionary data (instead of in-vivo data) for training. As such, even apart from eliminating the need for separate lengthy scans to obtain training data, it is expected that our approach is more versatile and generalizable, as covering the whole reasonable range of parameters is difficult to achieve while using in-vivo data for training.

### 2.5. Parameter estimation

As mentioned previously, a two-step parameter estimation approach is followed. As described earlier, in the first step, the GE data are used to obtain estimates of T1, T2* (and B1^+^), whereas in the second step, the SE data is used only to match for T2. For validation testing, the DNN-based parameter estimates were compared to those produced by the conventional method.

- *Conventional parameter estimation:* this is done using the magnitude of the MR signal and the pre-calculated dictionary using a maximum dot product approach. Based on a simplifying assumption that T2* can be represented by a mono-exponential decay (Rieger et al., 2017), both T2 and T2* can be estimated using the same dictionary, without the need to include T2 and T2* as different dimensions of the dictionary.
- *DNN-based parameter estimation:* as mentioned earlier, the input layer is fed the fingerprint time series for each voxel and the output layer produces B1^+^ corrected T1 and T2* estimates. To estimate T2 and based on these values, a mini-dictionary is constructed whereby only T2 varies between entries (here constituting 85 entries instead of the original 350,000 entries). Due to the use of this mini-dictionary, the simplest way to estimate T2 is by using conventional dictionary matching, which yielded the results almost instantaneously.

Note that the DNN-based T1 and B1^+^ estimates are fed into T2 estimation. This guarantees that as long as the T1 and B1^+^ estimates are accurate, errors in T2* estimation (also see Discussion) will not propagate into T2 estimation. Dictionary generation and data matching both were implemented in MATLAB R2019b (MathWorks, Natick, MA, USA) and C++ using custom-written codes.

### 2.6. Image acquisition

Imaging was performed on Siemens TIM Trio 3 T and Prisma 3 T systems (Erlangen, Germany) using a 32-channel receive-only head coil for the Trio and 64-channel head/neck coil for the Prisma. Specifically, the phantom and human validation scans were performed on the Trio, and the in vivo SMS-non-SMS comparison scans were performed on the Prisma. While this was driven by an institutional scanner upgrade, the Prisma system is nonetheless more appropriate for establishing the performance of state-of-the-art SMS.

### 2.7. Phantom validation

For validation, phantoms were built with varying aqueous concentrations of copper (II) sulfate (CuSO_4_) and agar (1-4% w/w). Seven small vials (15 mm in diameter) with different T1, T2, and T2* properties were placed inside a larger cylindrical plastic container filled with CuSO4 doped distilled water. Our MRF estimates were compared to those obtained from standard relaxometry techniques, referred to as “ground truth”.

Validation reference scans using “gold-standard” techniques were performed using the same spatial resolution as the single-band MRF acquisitions (1.1×1.1×3 mm^3^). To validate T1 estimations, inversion recovery turbo SE (IR-TSE) was used with eight different inversion times (TI = 50, 100, 200, 400, 800, 1600, 3200, 6400 ms), TR=10 s, TE=9.5 ms, BW/pixel=200 Hz and turbo-factor=8. For T2 validation, a single echo TSE sequence was repeated with seven echo times (TE=19, 38, 57, 76, 95, 114, 132, 152 ms), TR=5 s, BW/Pixel=200 Hz, Turbo-factor=8. T2* was measured using a multi-echo GRE with 12 echo times (TE=2, 5, 9, 13, 17, 25, 30, 40, 50, 60, 70, 80 ms) TR=1 s, BW/Pixel=390 Hz and FA=15°. B1^+^ maps were acquired using the double-angle method with a FLASH sequence, whereby FA1=60° and FA2=120°, TR/TE=5000/10 ms (Stollberger and Wach, 1996). Standard T1 fitting was done using complex data and the five-parameter model described by Barral et al. (Barral et al., 2010). A two-parameter model (S=M0 exp(TE/T2(*))) was used for both T2 and T2* by fitting a monoexponential curve to the data using a nonlinear (Levenberg-Marquardt) curve-fitting algorithm (MATLAB). Regions of interest (ROIs) were manually drawn around each of the small vials, excluding edge voxels. The mean and standard deviation of all the voxels in these ROIs were used for further comparisons.

### 2.8. In vivo validation

Following informed written consent, in accordance with the Institutional Research Ethics Board policy, in-vivo data from seven healthy human subjects (four males, mean±std: 27±6 years old) were acquired to complement the phantom validation. Three subjects were scanned on the Tim Trio scanner using only the single-band version of the sequence (1.1×1.1×3 mm^3^ voxels), and four subjects were scanned on the Prisma scanner with both single- and multi-band scans (1.7×1.7×3 mm^3^ voxels). Sequences and scanner information were similar to those described under “Phantom Validation” and “Pulse Sequence”. As in the phantom validation testing, conventional “gold-standard” scans were used to generate “ground-truth” parametric maps. In each subject, to avoid the prolonged acquisition time needed for “gold-standard” measurements (more than 30 minutes per slice for T1, T2, T2* and B1 validation scans), only one slice was acquired per subject for reference scans. Nevertheless, to probe the robustness of our technique, the slice position was varied across subjects to accommodate different levels of field homogeneity and anatomical properties. Furthermore, the performance of single-band and multi-band implementations compared to in-vivo was evaluated. Manual ROIs were drawn in several regions of the brain to represent grey matter (GM) and white matter (WM) regions as well as different field inhomogeneity scenarios. These ROIs were used for subsequent comparisons between sMRF and ground truth measurements.

### 2.9. Data and code availability

Under our institution’s ethics approval, the human data is not available for public sharing but the codes and phantom data are available upon request.

## 3. Results

As a demonstration of image quality, raw sMRF images from the phantom scan as well as one human scan are shown in Figures 4a and c. Compared to under-sampled spiral MRF images (Bipin Mehta et al., 2018), our images have visually higher SNR and minimal artifacts (in particular, no undersampling artifact). Also, despite a large matrix size (192×192), our use of GRAPPA acceleration with partial Fourier reconstruction appears successful in minimizing EPI-related image distortions. To demonstrate the quality of the single-voxel match between the dictionary entry and acquired data in our approach, we plot the conventional dictionary matched entry with our phantom and in-vivo data in Figures 4b and d, respectively, as a function of the frame number. Excellent correspondence between the sMRF data, the fitted dictionary element and the ground truth dictionary entry (simulated with Bloch equations using parameters determined by conventional relaxometry) can be seen, owing to the lack of the typically observed under-sampling artifact. As expected, the baseline signal level from the SE segment is lower than that from the GE segment, stemming from the lower baseline longitudinal magnetization that resulted from the continuous refocusing. Nonetheless, high overall T2 quantification accuracy is still achieved, likely since T2 is the only parameter estimated from this segment.

**Figure 4.**
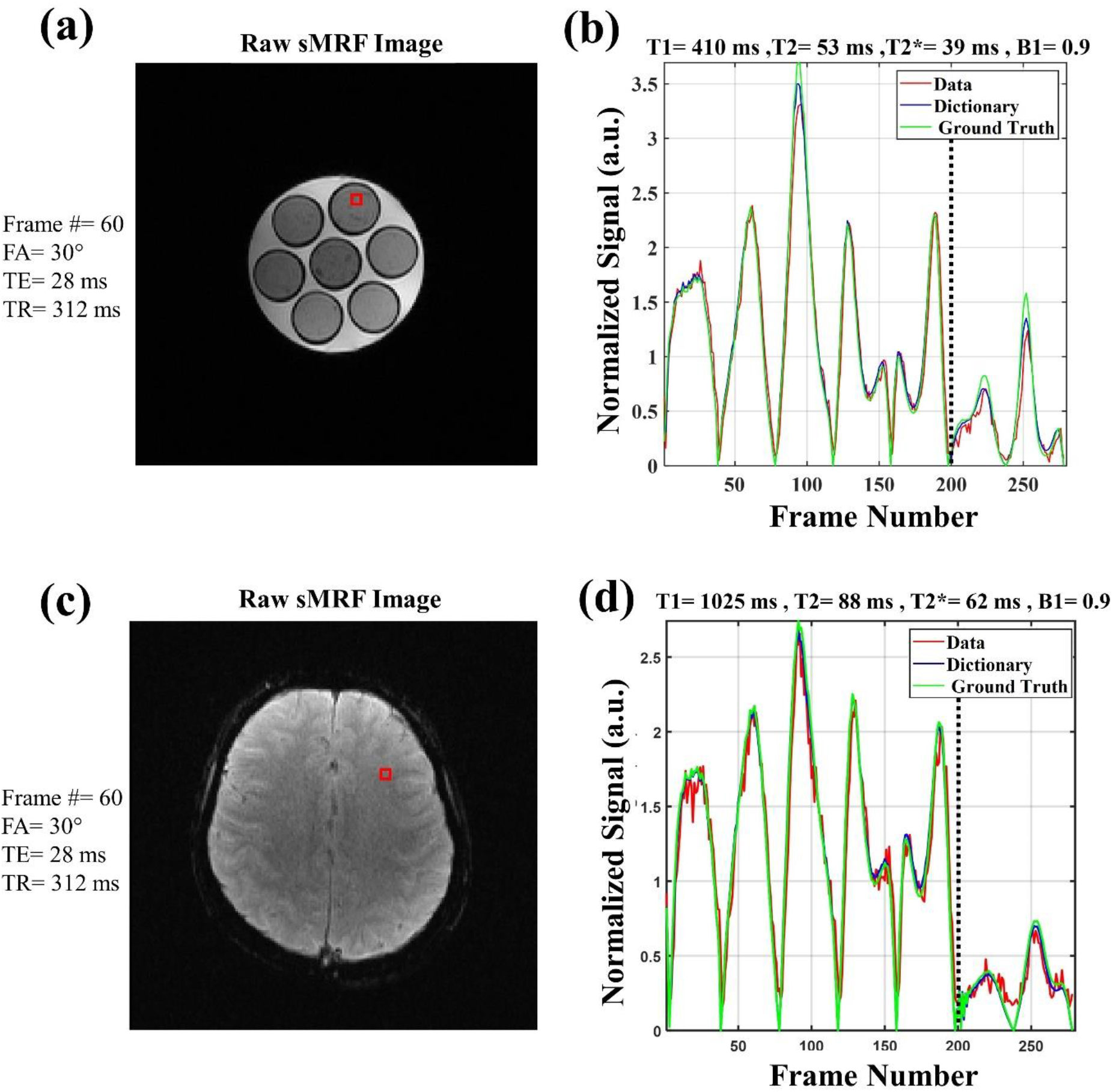
Quality of raw MRF images and the dictionary matching. (a) Raw sMRF image from the phantom scan (frame number=60, acquisition parameters listed in legend). (b) The quality of the match between the data, the selected dictionary entry and the ground truth dictionary element (created from ground-truth measurements), with the parameter estimates captioned on top for one voxel shown with a red square. (c) Raw sMRF image from the human scan (also frame number 60). (d) The quality of the match and the estimated values for one voxel shown with red square. In both cases, excellent agreement is found between the selected and the ground-truth dictionary entries. Dotted vertical lines in (b) and (d) show the start of the SE section.

The parametric maps obtained from phantom acquisitions using sMRF are shown alongside the “ground-truth” measurements (ground truth) in Figure 5. Different color schemes have been chosen here for each parameter for better distinction and visibility of different relaxation parameters, as was recommended and used previously (Obmann et al., 2019; Wang et al., 2018). Excellent correspondence between estimated and expected values (high R^2^ values) is achieved for all three relaxation parameters (T1, T2 and T2*). Particularly, for T2 quantification, which is one of the main novelties of our technique, the R^2^ is 0.996, the slope of the least square line is unity and the y-intercept is only 4 ms. For T1, the R^2^ is 0.993 and the slope of the least square line is 1.04. For T2*, R^2^ is 0.960 and the slope is 1.01.

**Figure 5.**
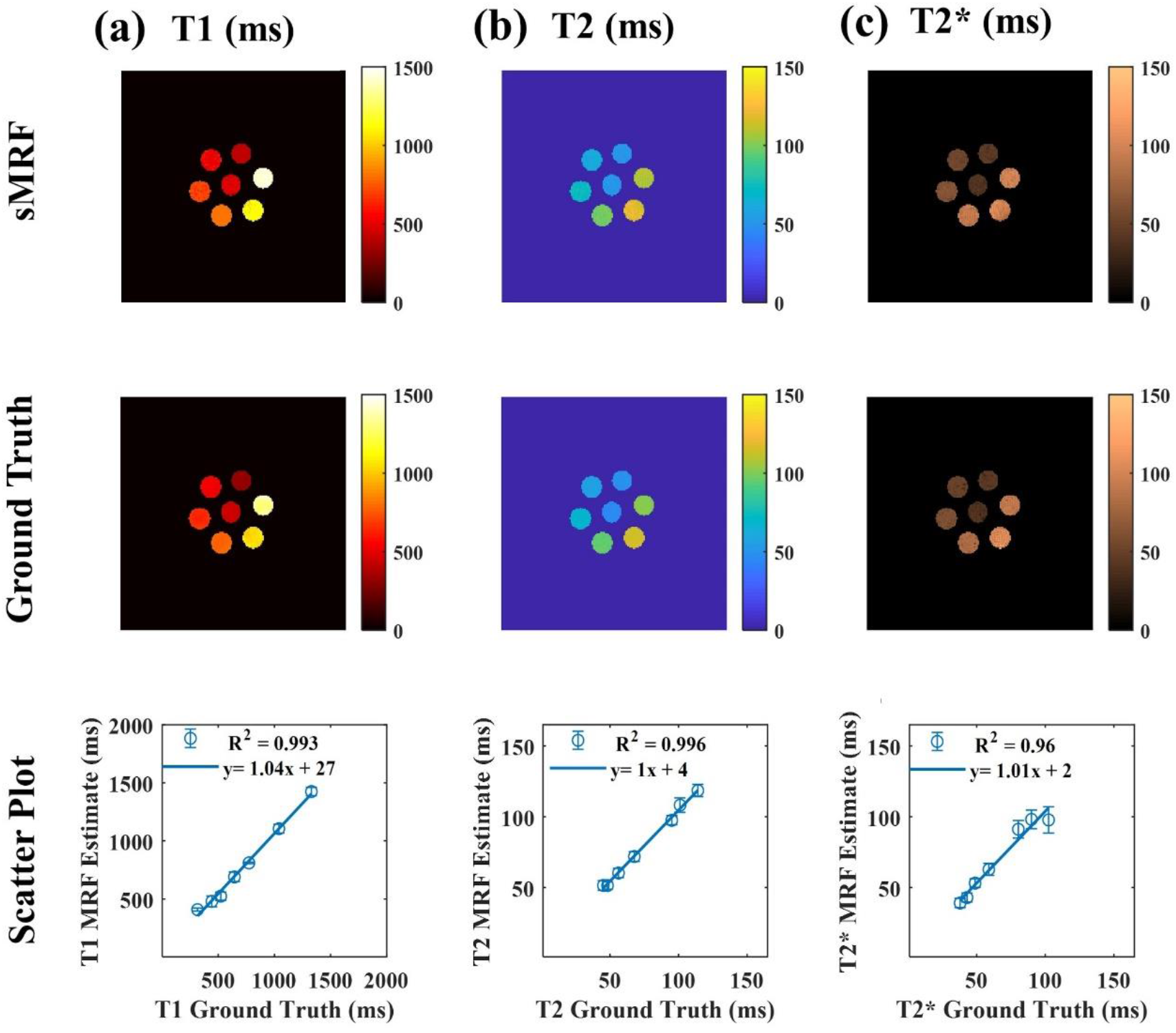
Phantom validation of sMRF approach based on gold-standard methods. (a,b,c) Strongly similar contrast was observed between our method and the validation scans that take ~30 min per slice. A high degree of quantitative agreement (R^2^ values and regression coefficients) in the parameter estimates is shown for all cases. Data are shown for the singleband implementation (acquisition time = 29 s per slice).

Validation of the single-band version of our in-vivo human sMRF results is shown in Figure 6, complete with three zoomed-in views of select regions. Their qualitative agreement with the standard measurements in terms of the image contrast is visible in this figure. As mentioned earlier, as the validation scans are lengthy, we acquired different slice positions, with each set of positions belonging to a different subject, resulting from our efforts to balance the number of slice positions and the length of scans per subject. Across all subjects and slice positions, similar contrast and range of estimated values can be observed. Moreover, good correspondence between our MRF-based B1^+^ maps and standard B1^+^ maps (double-angle method) is shown in Supplementary Figure 1 and is comparable with those shown in previous publications (Boudreau et al., 2017; Buonincontri and Sawiak, 2016; Körzdörfer et al., 2018). For voxels residing in the cerebral spinal fluid (CSF), a reduced agreement was observed between sMRF and ground-truth scans, which could be a result of pulsatile flow and the very long longitudinal relaxation time of CSF (see Discussion).

**Figure 6.**
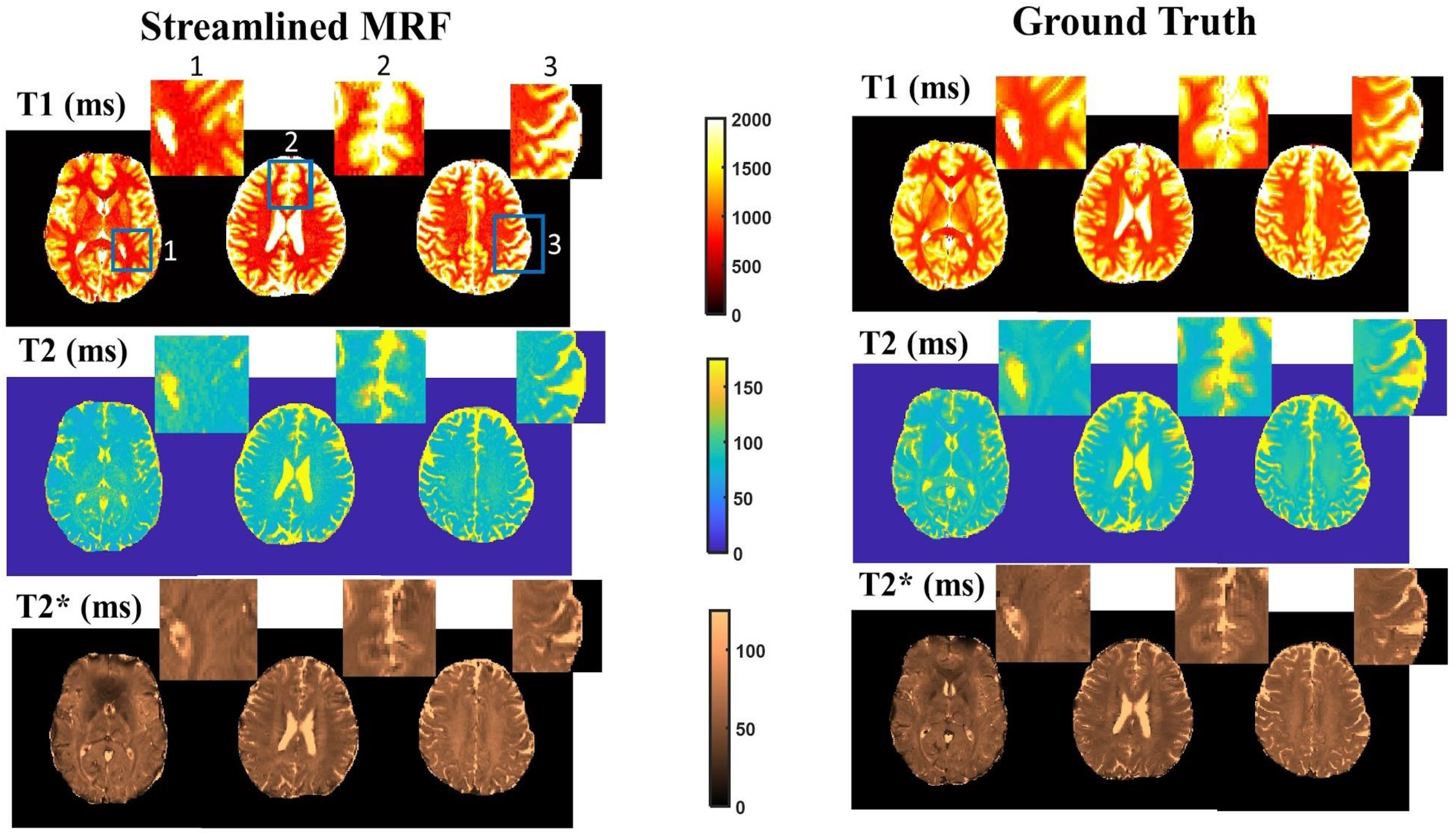
In-vivo validation of sMRF estimates and ground truth measurements for T1, T2 and T2* for different subjects and slice positions. Each slice is taken from a different subject and chosen in a different location to represent a range of field-heterogeneity conditions. The sMRF and ground truth results are in close agreement, except in some areas where cerebrospinal fluid (CSF) predominates

In Figure 7, we show the correspondence between sMRF estimates and ground-truth measurements for multiple selected ROIs in the slices shown in Figure 6, covering cortical and subcortical GM and WM. The values were taken from 3 different subjects, and averages taken for GM and WM separately. The means for both tissue types fall within the expected range and agree well with ground-truth measurements. Of particular interest is ROI 3 in Fig 7c. In this ROI, although the MRF T2* estimate is lower than the ground-truth value, likely as a result of through-plane field inhomogeneity (see Discussion), the T2 values seem to be unaffected, attesting to the decoupling between T2 and T2* estimates in our approach and the feasibility of estimating T2 using EPI-based MRF.

**Figure 7.**
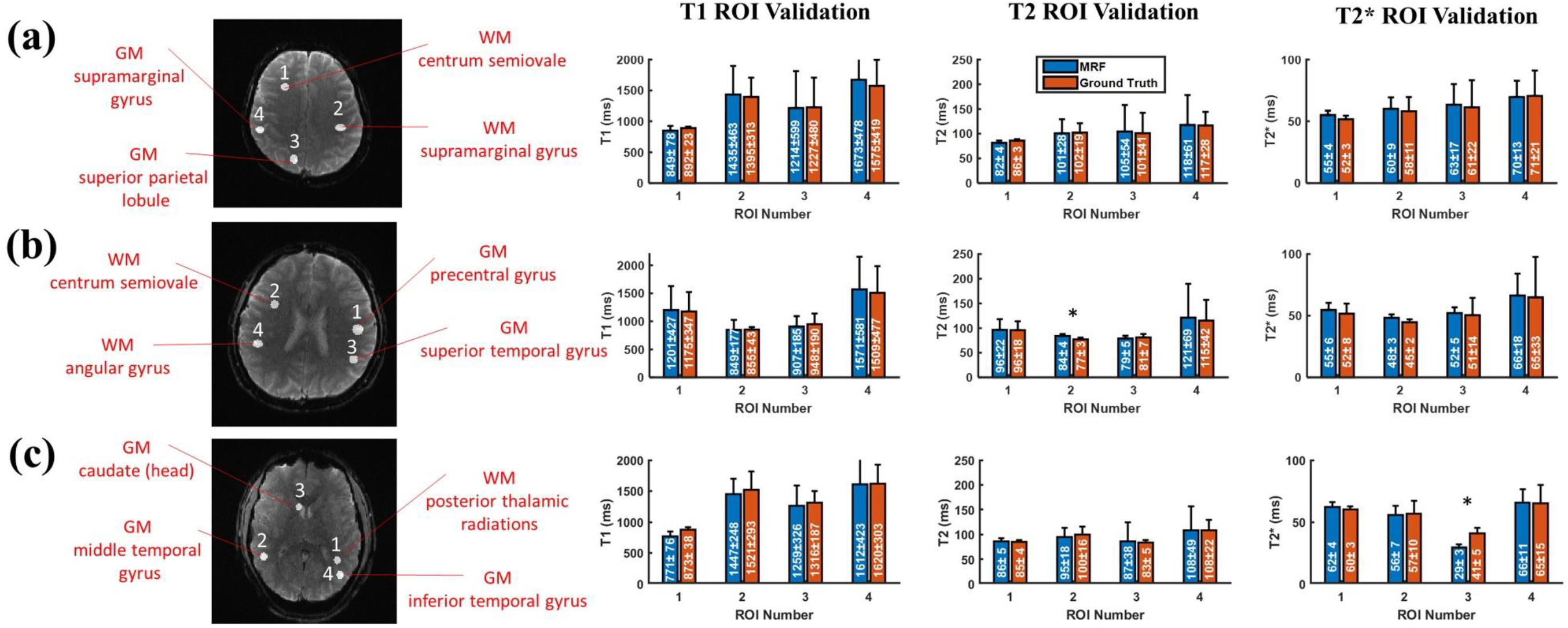
Correspondence between sMRF estimates and ground-truth measurements for multiple ROIs. Data are taken from 3 different subjects (one slice each). There is a good agreement between sMRF and ground truth estimates in different ROIs located in different GM and WM regions. The ROI means and standard deviations of the estimates are included in the bar graphs, with the error bars representing the standard deviation across the ROI. Asterisks indicate statistically different results (t-test, p<0.01), which is the case in only two cases (ROI 2 for T2 in (b) and ROI 3 for T2* in (c)). In this figure, the images are raw sMRF data from frame 60.

Both qualitative and quantitative comparison between the single-band and multiband (SMS-factor=4) versions of our method are shown in Figure 8. The comparison is shown visually for one subject as well as in the table for all the subjects and for two manually drawn ROIs in white and grey matter. Despite small differences, this figure indicates that comparable estimates can be obtained using either the single band or the multiband version of our sMRF method. Due to the rapid repetition of SMS pulses, especially in the form of refocusing pulses (the SE stage), we were sensitized to the specific absorption rate (SAR) levels. However, we found the SAR level reported by the scanner to be always well below the safety limits in all of our scans (at 16-19% of the maximum allowed SAR, depending on the subject).

**Figure 8.**
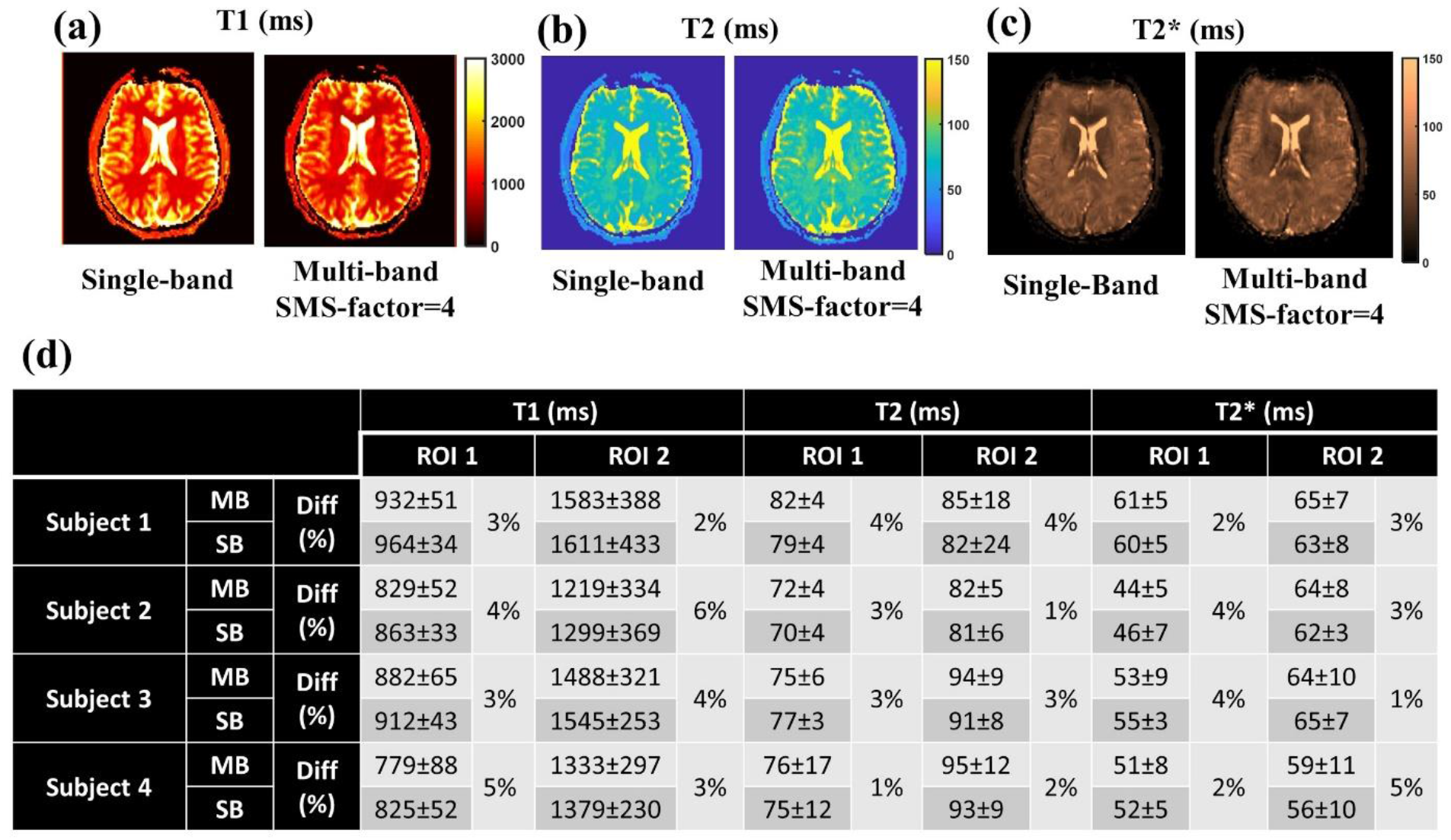
Comparison between the single-band (SB) and SMS (MB) versions of sMRF for multiple subjects. There is a good agreement between these two, both qualitatively, as can be seen in the parametric maps from one subject (a,b and c), and quantitatively, as can be seen in the table (d) summarizing the parameter estimates for white matter (ROI 1) and grey matter (ROI 2). The difference (“Diff’) between SB and MB estimates are shown as a percentage.

In Figure 9 we show the results of the DNN parameter estimation and its comparison with conventional dictionary matching. The quality of SMS acquisitions is demonstrated in slices with varying degrees of susceptibility effects (acquired using SMS option in the sequence). Parametric maps and difference maps are shown for one representative subject while the table shows the mean absolute differences (in percent) for all the subjects, which we assume to represent estimation errors by the DNN. The mean errors for T1, T2 and T2* are 7.4%, 3.6% and 6.0%, respectively, with a maximum error of 8% (in T1 for subject 2). Note that although simulated data alone was used for training the DNN here, the results compare well with a recent spiral DNN-MRF study (Fang et al., 2019a) which reported ~ 5% and 9% difference in T1 and T2 parameter estimates, respectively, in which in-vivo data was required for DNN training.

**Figure 9.**
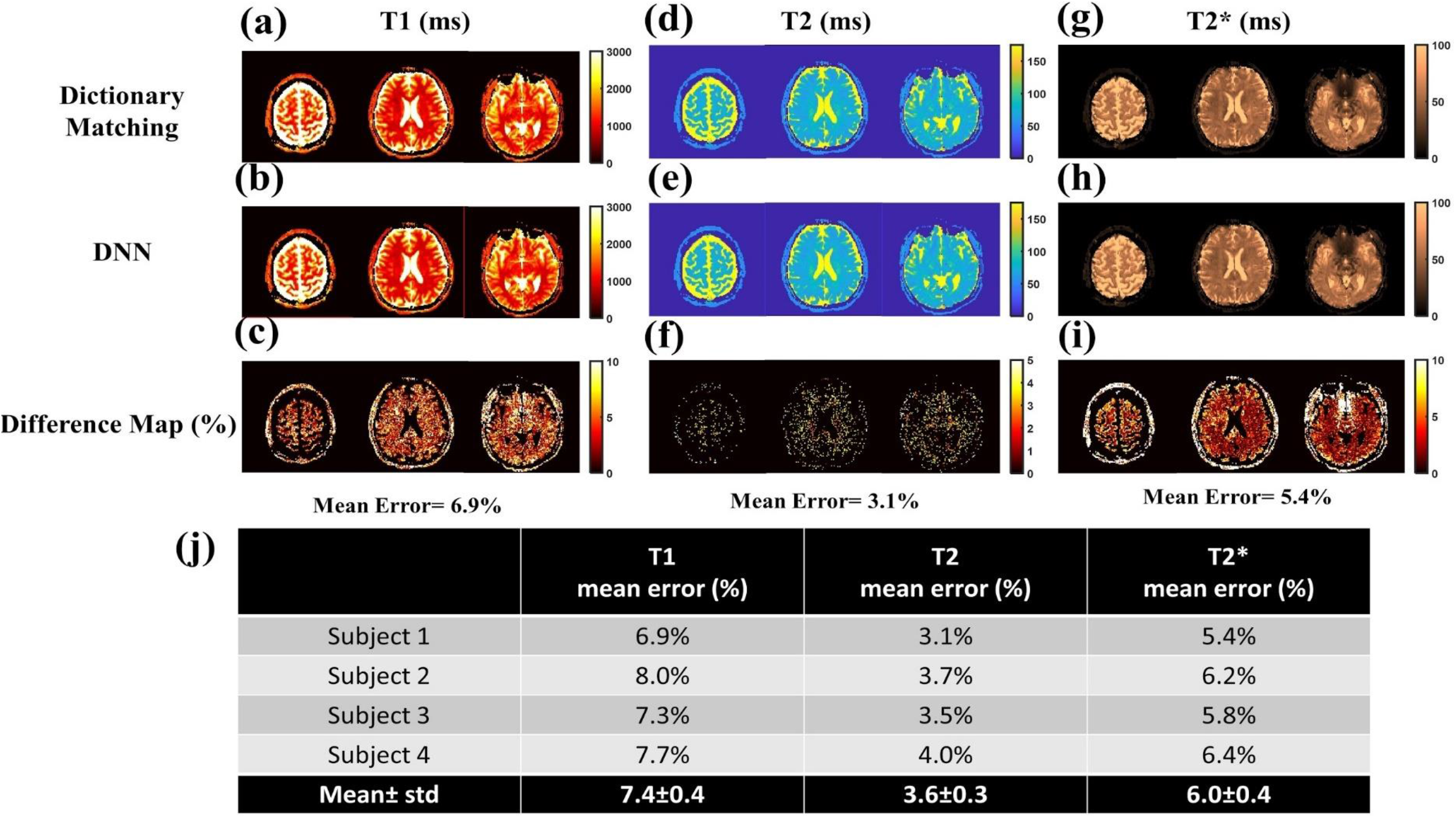
Comparison between DNN and the dictionary matching to obtain sMRF relaxation parameter estimates. The results are shown for 3 representative slices in one of the subjects (a-i). Moreover, a summary of the results for other subjects is provided (j) in terms of mean absolute error (%). The data presented were acquired using an SMS factor of 4 (voxel size = 1.7×1.7×3 mm^3^).

While running on a single PC (single-core 1.6 GHz, 16 GB of RAM) and using CPU, our DNN parameter estimation stage gave us estimates in almost 5 sec/slice, in stark contrast to around 5000 sec/slice in conventional dictionary matching; that is, the use of the DNN constitutes a ~ 1000 times acceleration. In terms of storage requirements, the size of the DNN was ~360 KB while the conventional dictionary was ~900 MB, so the DNN leads to a ~2500 times reduction in storage requirements. It should be noted that these exact values pertain to our own dictionary configuration, DNN architecture and hardware specifications, but similar factors of acceleration and storage savings should be expected in general

## 4. Discussion

In this work, an sMRF framework was introduced that extends and facilitates the application of MRF in multiple aspects. The following novelties contribute to the streamlining of MRF. First, T2 estimation was added to the previously introduced EPI-based MRF, making it possible to estimate all three main relaxation parameters in a single scan (T1, T2, and T2*). As also noted in previous work (Hermann et al., 2020; Rieger et al., 2018, 2017), EPI-based acquisitions make use of online instantaneous image reconstruction and produce images without noticeable under-sampling artifacts. Second, the utilization of simultaneous multi-slice (SMS) acceleration combined with slice interleaving makes this method capable of acquiring each slice in a few seconds, consequently enabling whole-brain coverage in just a few minutes (3 minutes here with SMS factor of 4). Thirdly, to reduce storage requirements and accelerate parameter estimation, a DNN was trained successfully, using simulated data alone, for in-vivo parameter estimation. It leads to orders of magnitude increase in speed of parameter estimation and similar degrees of reduction in storage requirements compared to when using conventional dictionary matching. In what follows, various aspects of the sMRF approach are discussed in more detail with respect to pertinent literature, and in the context of the results that were shown in Figs. 4-9.

### 4.1. EPI readout for MRF

The advantages of spiral imaging are well known and have been discussed earlier (Glover, 2012; Ma et al., 2013). It has been further shown that spiral MRF may provide benefits in terms of reduced estimation error as compared to Cartesian MRF while under-sampling is implemented (Stolk and Sbrizzi, 2019). Nonetheless, the practical feasibility of using an EPI readout for MRF is being increasingly recognized (Buonincontri and Sawiak, 2016; Rieger et al., 2017; Su et al., 2017), and will not be repeated here. As discussed previously, in this work, we capitalize on the fact that the EPI readout is readily available on commercial systems, which is complemented by fast online reconstruction with manufacturer-provided approaches for correcting gradient delays, imperfections, and nonlinearities. In addition, due to the absence of under-sampling artifacts in EPI images, we were able to use far fewer imaging frames for accurate parameter estimation than in the case of spiral MRF (Cohen and Rosen, 2017; Rieger et al., 2017). This leads to lower data sizes, faster dictionary generation (for DNN training), lower storage requirements and faster parameter estimation. As a case in point, our results show that the acquired signal in EPI-based MRF has high similarity with the matched noiseless dictionary entries generated by Bloch equation simulations (see Figure 4), owing to the absence of under-sampling artifacts. The high image quality of our approach facilitates the use of DNN parameter estimation (Cohen et al., 2018; Fang et al., 2019b). As the measured signals resemble the simulated ones, we were able to simplify the training process by not needing to acquire a large sample of undersampled data for realism of training. Furthermore, as motion correction has been recently raised as a means to improve MRF accuracy, the absence of undersampling artifacts in EPI images facilitates image-domain motion correction (Hermann et al., 2020). The high image quality also lends itself to possibly more accurate partial volume estimation through solving the inverse problem (Deshmane et al., 2019; Fang et al., 2019b).

### 4.2. EPI-based T2 estimation

For T1 and T2 MRF, spiral readouts are by far the most common, partly due to their ability to achieve very short TEs (Ma et al., 2013). However, in the case of EPI MRF, specifically, when using GE EPI, only T1 and T2* could be reliably estimated so far. Several groups have attempted to achieve T2 contrast by minimizing the TE (Cohen et al., 2018; Cohen and Rosen, 2017; Yang et al., 2019) as a way to suppress the T2* contamination. Yet, even with high in-plane acceleration factors and low partial Fourier factors, it is difficult to achieve an echo time of less than ~10ms for matrix sizes of 128×128 and above. Consequently, in these methods, T2* contamination is unavoidable and limited accuracy has been reported for T2 estimation, especially as compared to the case of T1 (Cohen and Rosen, 2017).

Here, we not only used a SE refocusing pulse to enhance T2 contrast but also merged this with a GE-EPI stage to afford T1, T2 and T2* estimation with B1^+^ correction using a single sequence. One may raise the alternative of implementing a single-stage SE-EPI MRF for T1/T2 estimation, but in reality, this option is much less promising. The main reason is the saturation of the longitudinal magnetization by the repeated refocusing pulses, which suppresses the signal, reduces the image SNR and leads to unreliable estimates (see Suppl. Figure. S2). This effect is conceptually similar to SNR loss in SE functional MRI while using very short TRs (Setsompop et al., 2012). Increasing the number of slices per TR (evenly distributed across the TR) is an intuitive way to increase the effective TR and consequently the baseline signal level, but it comes at the cost of losing a fraction of the T1 sensitivity due to the resultant sparser sampling of the inversion recovery curve. Our solution to this problem is to fit for T2 alone in the second stage of the sequence, a process that incorporates the T1 and B1^+^ information derived from the first (GE) stage. This approach is validated by our results, as the accuracy of our method for T2 estimation was found to be high (R^2^> 0.99, regression coefficient ~= 1, when compared to “ground-truth” measurements) in our phantom data as well as in in-vivo scans. The general principles and advantages of a multi-stage acquisition and fitting process are further discussed in a later section. Nonetheless, reduced accuracy in regions with abundant CSF was observed, likely due to pulsatile flow effects in the ventricles, but also due to the fact that our current approach is not optimized for covering extreme T2 values. Similar limitations for CSF-related parameter estimation have been found in other MRF studies as well (Bipin Mehta et al., 2018; Deshmane et al., 2019; Jiang et al., 2017; Ma et al., 2013), and can be partly addressed by extending the range of sensitivity in both acquisition and in dictionary construction (and consequently acquisition/matching times), if need be.

### 4.3. Comparison with alternative methods

Table 1 provides a systematic comparison of our method with selected existing qMRI approaches. Both MRF based and non-MRF based methods are covered in this table, with the former covered more comprehensively. The table gives a quick overview of where streamlined MRF (sMRF) stands. As shown in this table, no single method overtakes other methods in all aspects, and there are tradeoffs with all methods. sMRF is neither the highest in resolution nor the fastest in speed, but it combines ease of acquisition and estimation into a single method.

**Table 1.** Comparison of some existing multiparametric relaxometry techniques with MRF techniques, including the proposed streamlined MRF. This table highlights the persisting challenges in the rapidly growing field of qMRI and MRF, as well as the pros and cons of the incipient EPI-based MRF techniques. It should be noted that acquisition is not the only bottleneck in clinical MRF, as lengthy dictionary matching also has been cited as a main challenge.

| Method |  | Resolution (mm) | Scan time (sec/slice) | Online Image Recon | Readout type | T1 | T2 | T2* | B0 | B1 | PD |
| --- | --- | --- | --- | --- | --- | --- | --- | --- | --- | --- | --- |
| Non-MRF | DESPOT1 (Deoni et al. 2005) | 1 isotropic | ~ 6 | ✓ | Cartesian | ✓ | × | × | × | × | × |
|  | DESPOT2 (Deoni et al. 2005) | 1 isotropic | ~ 2 | ✓ | Cartesian | × | ✓ | × | × | × | × |
|  | IR-TrueFISP (Ehnes et al. 2013) | 1.5x1.5x10 | 5 | × | Radial | ✓ | ✓ | × | × | × | ✓ |
|  | TESS (Heule et al. 2014) | 1.2x1.2x4 | 28 | ✓ | Cartesian | ✓ | ✓ | × | ✓ | × | ✓ |
| MRF | MRF (Ma et al. 2013) | 2.3x2.3x5 | 12 | × | Spiral | ✓ | ✓ | × | ✓ | × | ✓ |
|  | PnP-MRF (Cloos et al. 2016) | 1.4x1.4x5 | 28 | × | Radial | ✓ | ✓ | × | × | ✓ | ✓ |
|  | Multi-slice EPI MRF (Rieger et al. 2018) | 1x1x3 | 7 | ✓ | EPI | ✓ | × | ✓ | × | × | ✓ |
|  | qRF-MRF (Wang et al. 2018) | 1.2x1.2x5 | 35 | × | Spiral | ✓ | ✓ | ✓ | ✓ | × | ✓ |
|  | Variable TE-MRF (Wyatt et al. 2018) | 1.1x1.1x5 | 19 | × | Spiral | ✓ | ✓ | ✓ | ✓ | × | × |
|  | FISP-SPGR MRF (Hong et al. 2018) | 1x1x3 | 40 | × | Spiral | ✓ | ✓ | ✓ | ✓ | × | ✓ |
|  | Streamlined MRF | 1.1x1.1x3<br>With MB=4:<br>1.7x1.7x3 | 29<br>With MB=4:<br>7 | ✓ | EPI | ✓ | ✓ | ✓ | × | ✓ | × |

Notably, there are three other published MRF papers that report simultaneous T1, T2 and T2* estimation using a spiral readout. In the work of Hong et al. (Hong et al., 2018), the acquisition time is 40 sec/slice for 1×1x3 mm^3^ resolution acquired over 800 frames in a multi-slice scheme (13 slices). In the work by Wang et al. (Wang et al., 2018), the acquisition time is 35 sec/slice for 1.2×1.2×5 mm^3^ resolution using 3000 frames in a single-slice acquisition scheme. Finally, Wyatt et al. (Wyatt et al., 2018) offer 19 sec/slice acquisition time for 1.1×1.1×5 mm^3^ resolution using 1000 shots (frames) in a single-slice acquisition scheme. In our work, our sMRF implementation calls for an acquisition time of 29 sec/slice for 1.1×1.1×3 resolution, and 7 sec/slice for 1.7×1.7×3 resolution with SMS-factor=4. However, our approach provides flexibility for the user to trade off spatial resolution for speed. Moreover, one advantage is that in both cases, we only needed to acquire 280 frames. Furthermore, sMRF also provides the possibility of using online image reconstruction and a DNN for parameter estimation that can be trained with simple Bloch simulations. As a result, we believe the sMRF pipeline is a strongly viable alternative to the spiral based approaches to simultaneously measure T1, T2, and T2*.

### 4.4. Multi-slice and SMS implementations

In the current implementation of our technique, multi-slice coverage comes with multiple tradeoffs. First, it increases the effective TR, thereby reducing the sampling rate of the T1 recovery curve. This may negatively impact T1 accuracy (Bipin Mehta et al., 2018), although our T1 estimation accuracy is high in reference to the gold-standard method (R^2^ = 0.993). On the other hand, the increase in effective TR increases the baseline signal, hence SNR, allowing for more accurate parameter estimations in general. Another tradeoff pertains to the baseline signal level in the SE-EPI stage. A short effective TR, while beneficial for T1 estimation, results in near saturation of the longitudinal magnetization, particularly due to the repeated application of refocusing pulses. This leads to low image SNR (see supporting information Supplementary Figure 2). Acquiring multiple slices, thereby increasing the effective TR to at least ~300 ms, alleviates this problem. Considering all these tradeoffs, in the single-band version of our sequence, 4-6 slices are deemed feasible. This is extended to the SMS case, in which the use of 4-6 slice groups would be optimal.

In our study, we did not notice any significant deleterious effect of imperfect slice profile, at our current slice gap of 50% (of the slice thickness). That is, our current implementation does not necessitate the customization of the commercially available RF pulse for improved pulse profile. Should smaller slice gaps ever be required, an alternative to pulse customization is to account for the slice profile effect in the dictionary.

Compared to the single-band case (resolution = 1.1×1.1×1.3 mm^3^), the reduced spatial resolution of the SMS version (1.7×1.7×1.3 mm^3^) stems from the limitations of GRAPPA and SMS when used in combination. Such limitations are well documented (Seidel et al., 2020; Setsompop et al., 2012), preventing us from using GRAPPA > 2 when high SMS acceleration is used concurrently. Nonetheless, if we reduced the SMS factor, at the expense of longer acquisition time, we would be able to achieve the resolution of 1.1×1.1×1.3 mm^3^, although in that case, multiple iterations of SMS would be needed for whole-brain coverage.

### 4.5. Two-stage approach

Sequences consisting of multiple stages with different sensitivities to specific parameters at each stage have been proposed in the MRF context in several previous instances (Buonincontri and Sawiak, 2016; Hong et al., 2018; Körzdörfer et al., 2018; Wang et al., 2018; Wyatt et al., 2018; Yang et al., 2019). Likewise, we use a two-stage approach to additionally estimate T2 (beyond T1 and T2*). Unique to our technique, as compared to other alternatives (such as using very short TEs (Yang et al., 2019)), the use of a refocusing pulse for T2 estimation guarantees better suppression of remaining T2* effects, as demonstrated by the high accuracy in the T2 estimates. Finally, In our approach, aside from the two-stage acquisition consisting of the GE and SE stages, the parameter estimation is also dual-stage, as the T2 estimation using the SE stage data incorporates T1 and B1^+^ information obtained from the GE section. By adding a second stage, the acquisition time increases by 40% (assuming 200 shots for the GE stage) compared to a single-stage scheme for T1/T2*/B1^+^ estimation. Thus, a main strength of a two-stage scheme comes from its high efficiency, in the sense that while a SE stage is indispensable (due to its T2 weighting), we use the T1 and B1^+^ values estimated from the first (GE) stage to reduce the complexity and data requirement for the T2 fitting. An associated strength of such an approach, particularly in the case of an EPI-based acquisition, is that we are able to, as proven by our results, prevent crosstalk between T2* and T2 estimates. These strengths may well generalize to a multi-stage design in which sensitivity to even more parameters is added in a semi-modular fashion. That is, certain stages may make use of estimates obtained from other stages to alleviate the challenge of the multi-parametric fitting.

### 4.6. Parameter estimate by deep learning

Deep learning has recently been suggested for parameter estimation in the MRF framework with success (Chen et al., 2019; Cohen et al., 2018; Fang et al., 2019a; Hoppe et al., 2017; Zhang et al., 2020). Deep learning-based parameter estimation allows much faster estimation and much lower storage requirements, as only the weights for the neurons (instead of the whole dictionary) need to be stored. As a result, the use of DNN removes the pattern-matching stage through direct estimation of parameters based on the input time series. However, as the under-sampled spiral data do not resemble simulated data, in vivo data or an estimation of empirical noise given the undersampling artifact are typically required for training (Virtue et al., 2018). This limits the quality of the training process or makes it more challenging, as the extent to which the DNN can represent different types of tissues and pathologies is limited by the availability of such experimental data. Without the training set that covers the whole parameter-space (Chen et al., 2019; Fang et al., 2019a), the resulting DNN-based estimates may understandably be limited in accuracy, which is a concern, especially in disease conditions.

In our method, due to the absence of significant artifacts, there is a strong similarity between Bloch simulations and the acquired data, as was also the case in EPI-based arterial-spin labeling MRF (Zhang et al., 2020). As such, we found a dictionary generated using Bloch simulations with just added Gusaaian noise sufficient for neural-network training, implying that we can generate potentially unlimited training data from a wide parameter space. Moreover, the use of DNN allows us to increase the dimensionality of our initial dictionary, thereby more thoroughly embodying the benefits of the MRF framework. Of course, it is understood that added sensitivity to extreme parameter values also entails modifications to the sequence.

Furthermore, our sMRF T2 estimation uses a simple dictionary matching process informed by DNN. This is because, having obtained the T1 and B1^+^ from the first stage, only a very small dictionary (mini-T2 dictionary) that covers the range of T2 values for a given T1 and B1^+^ is required. This dictionary is so small that the matching is almost instantaneous, precluding the need for a DNN at this stage. Of course, increasing dimensionality of the dictionary may necessitate the adoption of our DNN with expanded complexity.

### 4.7. Limitations and future work

While we showed strong evidence supporting the feasibility of using a streamlined dual-stage EPI-based MRF to rapidly estimate T2 in addition to T1 and T2*, we are mindful of the following limitations.

One limitation of our method is obviously the geometrical distortions due to the presence of B0 inhomogeneities that is common with EPI readouts. By using a combination of parallel imaging, partial Fourier acquisition and gradient and RF spoiling, we suppressed this effect to some extent, as shown in our results. Yet, it can still be a source of error in regions close to air cavities that suffer from high susceptibility differences (e.g. in Fig 6). For these reasons, some very recent qMRI studies that aimed to benefit from the efficiency of EPI have resorted to accelerated multi-shot approaches (Benjamin et al., 2019; Bilgic et al., 2019; F. Wang et al., 2019). Of course, this may come at the cost of more complex reconstruction and the need for phase navigator or data-driven approaches to deal with shot-to-shot motion.

Another limitation of our method comes from the representation of T2* as a mono-exponential decay (like T2), by assuming a Lorentzian distribution for the off-resonance values. The challenge arises in regions with high (mostly through-plane) field inhomogeneities, in which this assumption may not hold. Figures 5 and 6 show examples of this scenario, in which our MRF approach underestimates T2* in the frontal lobe. In such cases, aside from explicitly estimating B0 (Hong et al., 2018; C. Y. Wang et al., 2019), a reduction of slice thickness and more advanced iterative shimming or z-shimming (Nam et al., 2012; Yang et al., 1998) schemes can be considered as possible solutions.

Moreover, while we have mostly shown the applicability and accuracy of our approach in a relevant range for brain tissue T1, T2 and T2* estimation, values that are far above and below this range can present challenges. In particular, to accurately estimate very short T2 (<30 ms), we should shorten the minimum TE in the SE stage using higher acceleration factors or partial Fourier reductions. Also, estimating very short T1 (~<300 ms) is equally challenging due to our sampling rate of the T1-decay curve. These attributes will be investigated in our future work.

Finally, a more ambitious but important goal is to eventually optimize the pattern of TR, TE and FA such that our approach can be sped up (with the number of frames reduced) without sacrificing accuracy. In this case, several metrics need to be accounted for, including the baseline signal level (which affects image and temporal-SNR), the separability of the dictionary elements based on temporal and spatial noise distributions, slice-profile imperfections, and magnetization transfer effects, to name a few. Hardware limitations are also likely to sway the optimization for each scanner type. Therefore, finding a globally optimized pattern for EPI MRF (or MRF in general) is non-trivial and outside the scope of the current work.

## 5. Conclusion

In this work, we present a streamlined MRF method (sMRF) to estimate T2, T1 and T2* simultaneously with B1^+^ correction. The use of an EPI readout provides ease of implementation of this approach as well as allows the adoption of simultaneous multi-slice acceleration for fast whole-brain coverage. Furthermore, a deep-learning based matching stage that can be trained with simple Bloch simulations allows estimation of parameters in only a few seconds per slice, with low percent differences compared to conventional dictionary matching. Overall, this approach introduces a novel framework that merges a number of novel techniques to pave the way for the broader adaptation of MRF.

## Appendix

To estimate the minimum number of frames required for the SE-EPI segment, we first obtained a rough estimate of the data SNR. Then, the number of required SE frames was determined through Monte Carlo simulations. We generated a small dictionary with T1 ranging from 500 ms to 1500 ms and T2 ranging from 50 ms to 150 ms (~1200 entries) while assuming homogeneous B1^+^. Subsequently, Gaussian white noise (based on the estimated experimental SNR) was added to the dictionary. Pattern matching was done and the mean relative error of T2 estimation was calculated to investigate the relationship between T2 error and the number of acquired SE frames. Based on this, we found that 80 SE frames leads to less than 2% error in T2 estimation and seems a reasonable choice by balancing acquisition time and accuracy (See Suppl. Figure S3).

## Declarations of interest

None.

## Acknowledgments

We are grateful to Drs. Kamil Uludag, Jean Chen, David Norris, and Peter Koopmans for their helpful comments during the preparation of the manuscript. We also thank Baycrest Hospital and Dr. Jean Chen for partial financial support. We also thank Fatemeh Rastegar and Dr. Ata Jodeiri for their help.

---

**Suppl. Figure S1.**
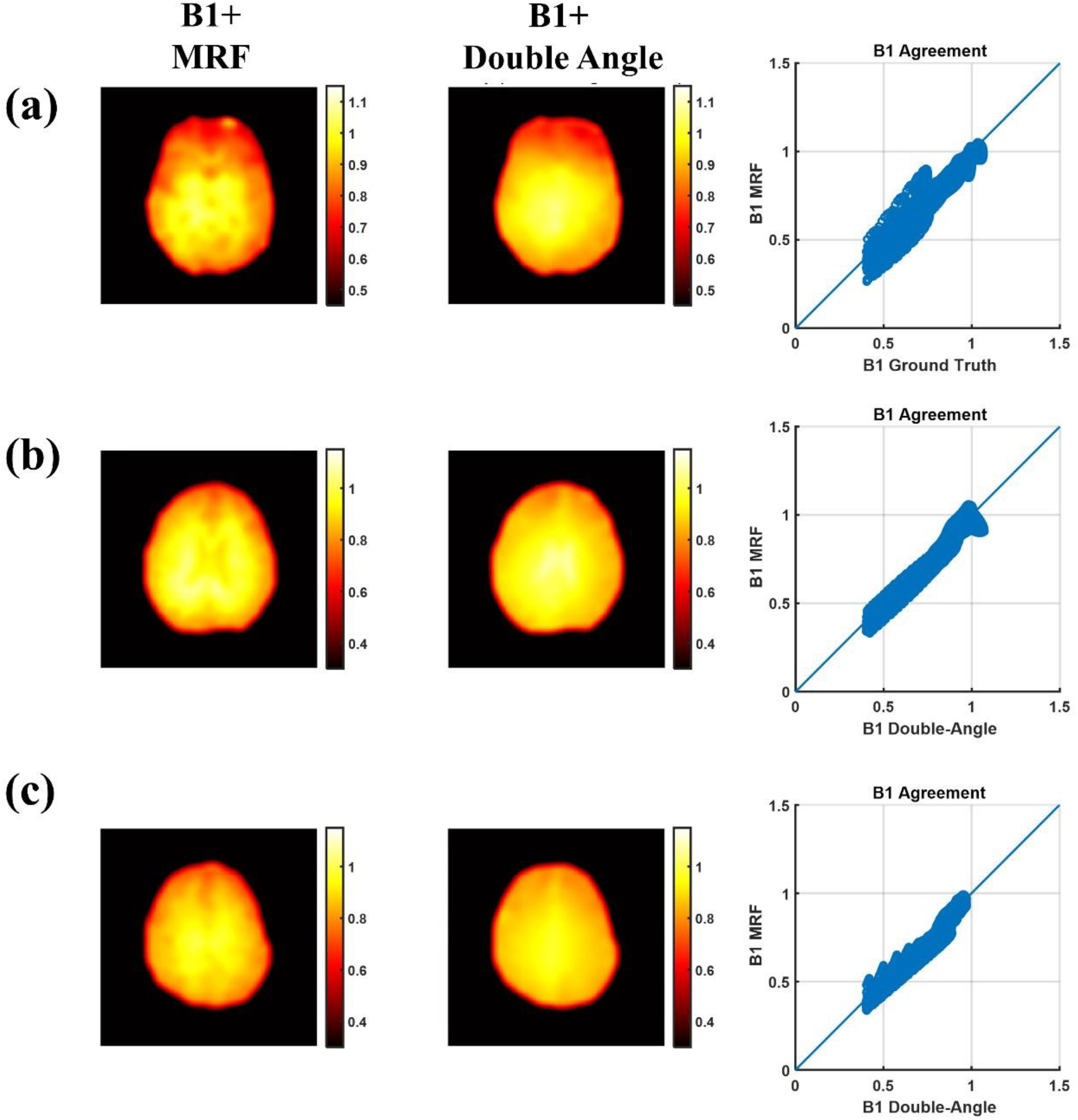
Correspondence between MRF B1^+^ maps and the ground truth method (double angle) for the 3 slices shown in Fig 6. There is a reasonable agreement between these two methods as is evident from the visual comparison and also the scatter plots. Scatter plots contain all the voxels in the represented slice. Gaussian smoothing has been applied as is standard in the B1 mapping literature based on (Boudreau et al., 2017).

**Suppl. Figure S2.**
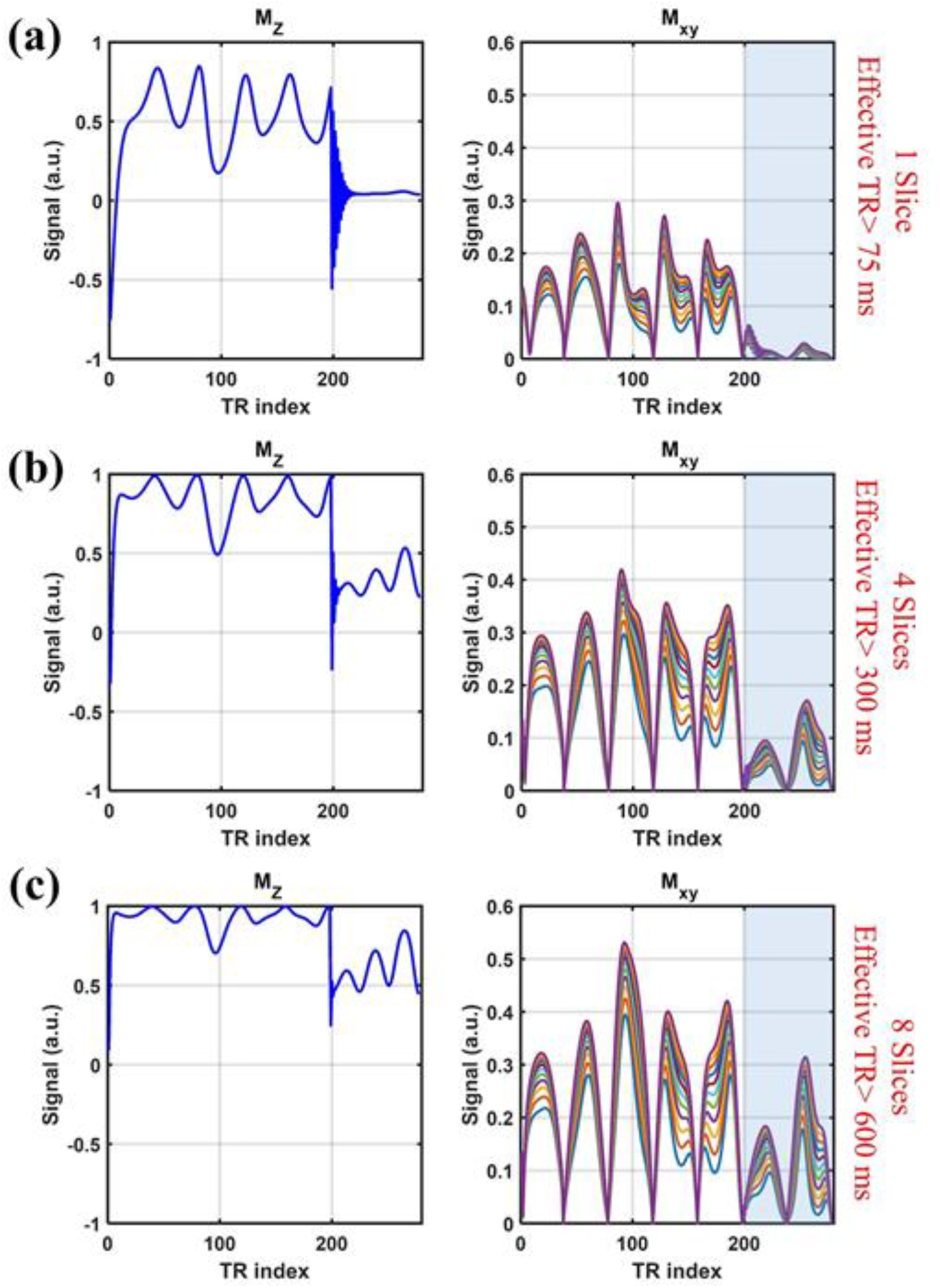
Exploring the effect of increasing longitudinal-recovery time on the baseline signal level. In each case, the recovery time is controlled by varying the effective TR by the number of slices acquired per TR. The relationship between the longitudinal/transverse magnetization and the recovery time is explored using Bloch simulations for a representative tissue with T1 of 800 ms and T2 values ranging between 50-150 ms. (a) single slice per TR with recovery time > 75 ms. (b) 4 slices per TR with recovery time > 300 ms. (c) 8 slices per TR with a recovery time > 600 ms. The shaded region represents the SE segment. Increasing the recovery time clearly leads to increased baseline signal in the SE segment (as well as the GE section).

**Suppl. Figure 3.**
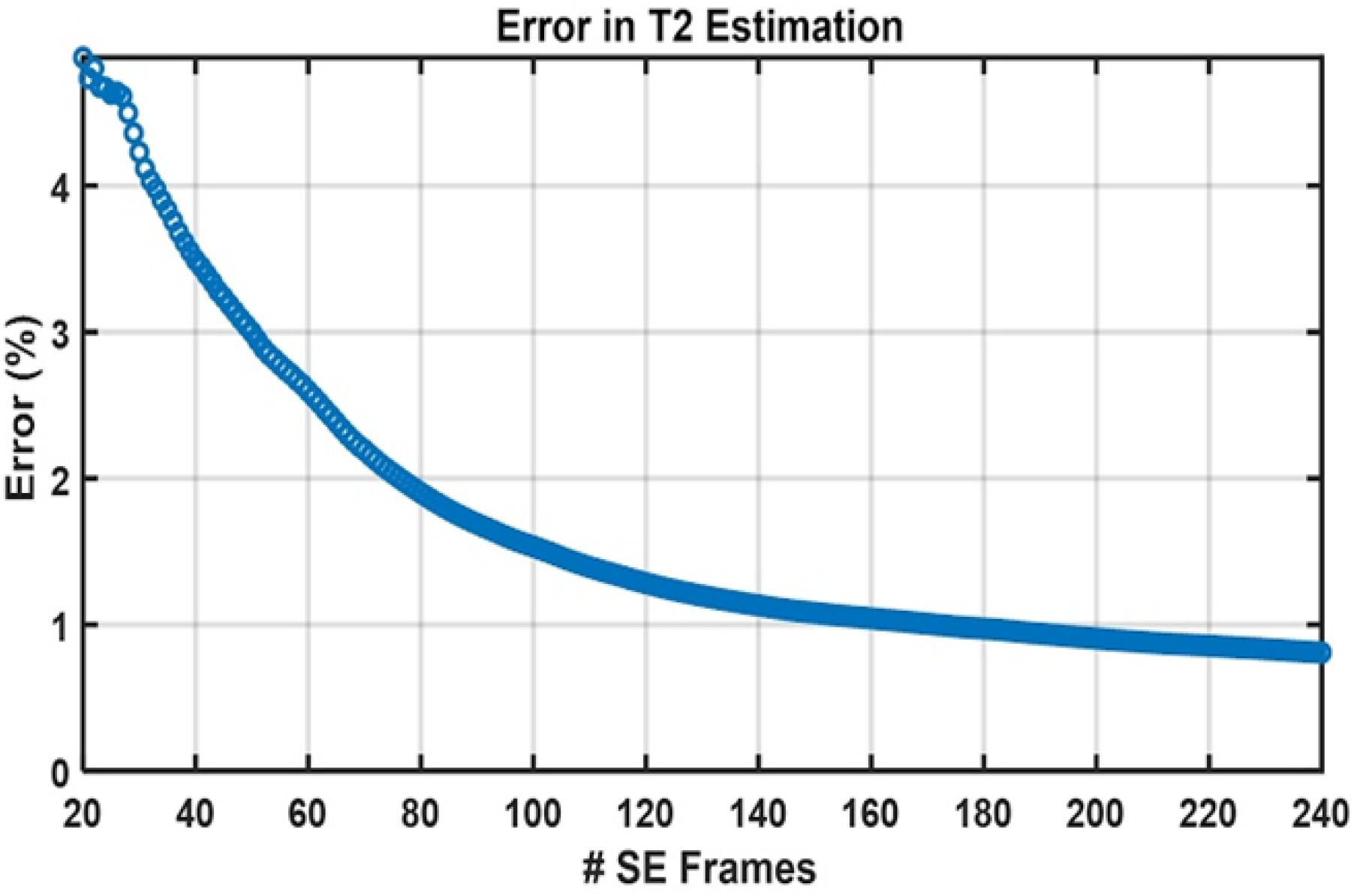
Error (%) in T2 estimation based on the number of frames used for the SE segment. The knee of the curve is at approximately 80 frames, which was used as a reasonable compromise between acquisition time and accuracy.

## Notes

### Competing Interest Statement

The authors have declared no competing interest.

